# White matter microstructural and morphometric alterations in autism: Implications for intellectual capabilities

**DOI:** 10.1101/2021.10.11.464005

**Authors:** Chun-Hung Yeh, Rung-Yu Tseng, Hsing-Chang Ni, Luca Cocchi, Jung-Chi Chang, Mei-Yun Hsu, En-Nien Tu, Yu-Yu Wu, Tai-Li Chou, Susan Shur-Fen Gau, Hsiang-Yuan Lin

**Affiliations:** Institute for Radiological Research, Chang Gung University and Chang Gung Memorial Hospital, Taoyuan, Taiwan; Department of Psychiatry, Chang Gung Memorial Hospital at Linkou, Taoyuan, Taiwan; Clinical Brain Networks Group, QIMR Berghofer Medical Research Institute, Brisbane, Queensland, Australia; Department of Psychiatry, National Taiwan University Hospital and College of Medicine, Taipei, Taiwan; YuNing Clinic, Taipei, Taiwan; Department of Psychiatry, University of Oxford, Oxford, United Kingdom; Department of Psychiatry, Keelung Chang Gung Memorial Hospital, Keelung, Taiwan; Department of Psychology, National Taiwan University, Taipei, Taiwan; Azrieli Adult Neurodevelopmental Centre, Campbell Family Mental Health Research Institute, and Adult Neurodevelopmental and Geriatric Psychiatry Division, Centre for Addiction and Mental Health, Toronto, Ontario, Canada; Department of Psychiatry and Institute of Medical Science, Temerty Faculty of Medicine, University of Toronto, Toronto, Ontario, Canada

**Keywords:** Autism spectrum disorder, fixel-based analysis, intellectual disabilities, minimally verbal status, cerebellum, diffusion MRI

## Abstract

**Background:** Neuroimage literature of autism spectrum disorder (ASD) has a moderate-to-high risk of bias, partially because those combined with intellectual impairment (II) and/or minimally verbal (MV) status are generally ignored. We aimed to provide more comprehensive insights into white matter alterations of ASD, inclusive of individuals with II (ASD-II-Only) or MV expression (ASD-MV).

**Methods:** Sixty-five participants with ASD (ASD-Whole; 16.6±5.9 years; comprising 34 intellectually able youth, ASD-IA, and 31 intellectually impaired youth, ASD-II, including 24 ASD-II-Only plus 7 ASD-MV) and 38 demographic-matched typically developing controls (TDC; 17.3±5.6 years) were scanned in accelerated diffusion-weighted MRI. Fixel-based analysis was undertaken to investigate the categorical differences in fiber density (FD), fiber cross-section (FC), and a combined index (FDC), and brain-symptom/cognition associations.

**Results:** ASD-Whole had reduced FD in the anterior and posterior corpus callosum and left cerebellum Crus I, and smaller FDC in right cerebellum Crus II, compared to TDC. ASD-II, relative to TDC, showed almost identical alterations to those from ASD-Whole vs. TDC. ASD-II-Only had greater FD/FDC in the isthmus-splenium of callosum than ASD-MV. Autistic severity negatively correlated with FC in right Crus I. Non-verbal full-scale IQ positively correlated with FC/FDC in cerebellum VI. FD/FDC of the right dorsolateral prefrontal cortex showed a diagnosis-by-executive function interaction.

**Limitations:** We could not preclude the potential effects of age and sex from the ASD cohort, although statistical tests suggested that these factors were not influential. Our results could be confounded by variable psychiatric comorbidities and psychotropic medication uses in our ASD participants recruited from outpatient clinics, which is nevertheless closer to a real-world presentation of ASD. The outcomes related to ASD-MV were considered preliminaries due to the small sample size within this subgroup.

Finally, our study design did not include intellectual impairment-only participants without ASD to disentangle the mixture of autistic and intellectual symptoms.

**Conclusions:** ASD-associated white matter alterations appear driven by individuals with II and potentially further by MV. Results suggest that changes in the corpus callosum and cerebellum are key for psychopathology and cognition associated with ASD. Our work highlights an essential to include understudied sub-populations on the spectrum in research.

## INTRODUCTION

Autism Spectrum Disorder (ASD) is a neurodevelopmental condition, characterized by struggling in socio-communication and the restricted/repetitive behaviors and interests [1]. ASD is associated with categorically and dimensionally neurodevelopmental alterations in brain structures and function, contributing to suboptimal information processing underpinning social-communication, sensorimotor integration, and executive control processes [2, 3].

Supporting this observation and extending this understanding to the level of brain connections, studies using diffusion magnetic resonance imaging (dMRI) showed that altered white matter (WM) microstructural properties associated with ASD exist in WM tracts interconnecting brain regions/systems involved in social processing and executive control [4–7]. For example, increased mean diffusivity (MD) or decreased fractional anisotropy (FA) in the corpus callosum (CC), uncinate fasciculus, superior longitudinal fasciculus, and frontal and temporal thalamic projections have been reported in people with ASD, compared to typically developing control (TDC). However, high variability of findings is commonly noted across studies, partially because of limitations and factors that can affect dMRI results. Specifically, voxel-based analysis (VBA) and tractography of dMRI popularly involve the analysis based on the diffusion tensor model, i.e., DTI [8]. But this model fails to resolve the multiple fiber orientations in regions containing crossing fibers, which exists in most WM voxels [9]. Thus, advanced dMRI models to address this methodological challenge are crucial to advance understanding of the WM pathology associated with ASD.

The state-of-the-art fixel-based analysis, or FBA [10], provides the fiber tract-specific analysis to estimate the quantitative metrics associated with a single fiber population within a voxel (called ‘fixel’), as opposed to analyses of voxel-averaged metrics. FBA has been shown to be more sensitive and interpretable than voxel-wise methods [11, 12], in terms of better reflection of the local WM features and morphology, by providing quantitative information of each fiber bundle cross-section given multiple crossing fibers within a voxel. Recently, the FBA has been applied to dMRI studies on ASD [13, 14], showing promising but mixed results. Specifically, Dimond *et al.* [13] reported that able youth with ASD have lower microstructural density in the right uncinate and arcuate fasciculi alongside CC-splenium whose fixel fiber density (FD; defined in Methods) is associated with social impairment. Kirkovski *et al.* [14] nevertheless only observed altered micro-/macro-structure in the posterior midbody of CC in able female adults with ASD. This inconsistency may, in part, reflect different sampling protocols of dMRI (e.g., number of gradient directions and b-values, resulting in variations of diffusion signals [15] across studies, as well as heterogeneity among individuals with ASD. Moreover, both studies only focused on individuals with relatively intact language and cognitive functioning, limiting the representativeness of the autistic cohort.

In addition to the above methodological considerations, around 50% of children with ASD have cognitive and intellectual impairments (i.e., IQ scores <85, representing intellectual disabilities (ID) plus borderline intelligence [16]), and about 30% of children with ASD remain minimally verbal (MV) by reaching their school age [17]. Unfortunately, these “low-functioning” individuals are understudied and often excluded from neuroimaging research. Despite prominent heterogeneity in designs, samples and findings, results from the limited existing studies suggest that people with ASD and ID have alterations of gray matter (GM) and WM morphology in diffuse regions implicated in ASD pathology [18]. Specifically, children with MV ASD showed structural disruptions in language pathways [17, 18]. In the Autism Phenome Project (APP) or Girls with Autism: Imaging of Neurodevelopment (GAIN) cohort [19, 20], children with ASD across the spectrum of developmental levels showed a transition from increased (during toddlerhood) to decreased (2 years later) FA in the CC, superior longitudinal fasciculus, cingulum, and internal capsule. The paucity of studies including individuals with co-occurring developmental disabilities [18] do, however, bias the understanding of brain bases of ASD [21].

Using the state-of-the-art dMRI acquisition (four b-values with ∼200 gradient directions), data processing framework (FBA), as well as the extensive psychopathological measures, the current study aimed to provide more comprehensive insights into the structural brain changes underpinning ASD. The current study included individuals with ID and MV, who are generally left out in the current lore. We hypothesized that the FBA could highlight potential differences in ASD that are driven by intellectual challenges. Based on previous work, we expected that intellectually able individuals with ASD would show alterations of WM fibers in the CC [13, 14], uncinate and superior longitudinal fasciculi, and the thalamic radiation. Individuals with ASD plus cognitive impairment (including those with MV expression) would show alterations with broader spatial involvement. Autistic traits, cognitive impairment, and poor adaptive function were in fact expected to map onto fixel pathology in the tract interconnecting social and cognitive brain networks.

## METHODS

### Procedures and Participants

The study was approved by the Research Ethics Committee of National Taiwan University Hospital (#201512238RINC). Participants with ASD (aged 8-30 years) were recruited from the outpatient psychiatric clinic at National Taiwan University Hospital, Chang Gung Memorial Hospital, Linkou, and Yuning Clinic. Age-matched TDC were recruited from neighborhoods with a similar social-economic environment to the ASD group. Before implementing the experiment, for those capable of giving consent themselves (i.e., showing capacity to understand the protocol), written informed consent was obtained from each participant and their parents. For those who were incapable of consenting, assent was sought and their substitute decision-maker signed the informed consent.

Eighty-six participants with a clinical diagnosis of ASD, made by child psychiatrists based on the DSM-5, and 39 TDC initially joined the study. The ASD diagnosis in all participants in the ASD group were further confirmed by the Autism Diagnostic Observation Schedule, ADOS [22, 23] and Autism Diagnostic Interview-Revised, ADI-R [24, 25].

Parents of all participants received an interview by the senior author (H.-Y.L.) using the Kiddie-Schedule for Affective Disorders and Schizophrenia-Epidemiological Version (K-SADS-E) for DSM-5 [26], to evaluate co-occurring psychiatric disorders in patients and to ensure that TDC were free of any mental health issues. Exclusion criteria for all participants in the ASD group included: Any acute or unstable medical illness; history of psychosurgery or head trauma; any active grand mal seizures in the past one year; known genetic causes contributing to ASD or ID; a history of bipolar, psychotic or substance use disorders; current suicidal ideation; pregnancy.

In addition to clinical assessments, all participants received the intellectual assessments using the Wechsler Intelligence Scale-4^th^ edition, either WAIS-IV [27] or WISC-IV [28] with the age cutoff of 16 years, and Leiter International Performance Scale-Revised, Leiter-R [29]. Their parents completed several scales to estimate participants’ behaviors and function, including the Social Responsiveness Scales, SRS [24], for autistic traits, the Vineland Adaptive Behavior Scales, VABS [30, 31], for adaptive function (adaptive behavior composite), and the Behavior Rating Inventory of Executive Function, BRIEF [32], for daily-life executive function.

After the quality control steps detailed as follows, 65 participants with ASD (6 females, age: 16.6±5.9 years) and 38 TDC (8 females, age: 17.3±5.6 years) were included in the final dMRI analysis (Table 1). Among those excluded, 14 participants with ASD failed to complete the dMRI scan; 7 participants with ASD and 2 TDC exhibited excessive motion during the dMRI scan.

**Table 1.** Demographic data and clinical features of participants

|  | Typically Developing Control (TDC)<br>n=38 |  | Intellectually Able ASD (ASD-IA)<br>n=34 |  | Intellectually Impaired ASD (ASD-II)<br>n=31 |  | P-value | Post-hoc test |
| --- | --- | --- | --- | --- | --- | --- | --- | --- |
|  | Mean | SD | Mean | SD | Mean | SD |  |  |
| Sex (M:F) | 30:8 |  | 30:4 |  | 29:2 |  | 0.20<sup>a</sup> | - |
| Age (years) | 17.3 | 5.65 | 15.9 | 5.46 | 17.3 | 6.41 | 0.55<sup>b</sup> | - |
| RMS (mm) | 0.399 | 0.197 | 0.407 | 0.184 | 0.463 | 0.216 | 0.40<sup>c</sup> | - |
| Medication (Y:N) | 0:38 |  | 14:20 |  | 9:22 |  | 0.31<sup>a</sup> | -<sup>d</sup> |
| Comorbidity (Y:N) | 0:38 |  | 26:8 |  | 24:7 |  | 0.93<sup>a</sup> | -<sup>d</sup> |
| ICV (cm<sup>3</sup>) | 1604.7 | 142.1 | 1588.8 | 114.5 | 1626.2 | 104.7 | 0.47<sup>b</sup> | - |
| *Cognition and Symptoms* |  |  |  |  |  |  |  |  |
| FSIQ (WISC/WAIS-IV) | 112.2 | 11.7 | 104.5 | 14.2 | 65.7 | 13.0 | <0.001<sup>c</sup> | ASD-II < ASD-IA, TDC |
| NVFIQ (Leiter-R) | 121.6 | 10.2 | 115.0 | 18.3 | 77.6 | 26.3 | <0.001<sup>c</sup> | ASD-II < ASD-IA, TDC |
| VABS-ABC | 113.5 | 15.3 | 89.2 | 15.1 | 66.7 | 9.6 | <0.001<sup>c</sup> | ASD-II < ASD-IA < TDC |
| BRIEF-GEC | 92.3 | 19.1 | 155.3 | 28.5 | 152.5 | 25.5 | <0.001<sup>c</sup> | ASD-II, ASD-IA > TDC |
| SRS-Total | 18.0 | 10.4 | 87.6 | 27.0 | 93.8 | 23.3 | <0.001<sup>c</sup> | ASD-II > ASD-IA > TDC |
| ADOS-2 CSS | N/A | N/A | 5.0 | 2.1 | 6.8 | 2.1 | 0.002<sup>c</sup> | ASD-II > ASD-IA<sup>d</sup> |

<sup>a</sup>: Chi-square; <sup>b</sup>: ANOVA; <sup>c</sup>: Kruskal-Wallis; <sup>d</sup>: comparison between ASD-IA and ASD-II
Acronyms: RMS: relative Root-Mean-Square of framewise displacement during diffusion MRI; ICV: Intra-Cranial Volume; FSIQ: Full-Scale Intelligence Quotient; WISC-IV: Wechsler Intelligence Scale for Children-4th edition; WAIS-IV: Wechsler Adult Intelligence Scale-4th edition; NVFIQ: Non-Verbal Intelligence Quotient; Leiter-R: Leiter International Performance Scale-Revised; VABS-ABC: Vineland Adaptive Behavior Scales Adaptive Behavior Composite; BRIEF-GEC: Behavior Rating Inventory of Executive Function Global Executive Composite; SRS-Total: Social Responsiveness Scales Total Raw Score; ADOS-2 CSS: Autism Diagnostic Observation Schedule 2 Calibrated Severity Score.

The whole ASD group (hereinafter ASD-Whole) was categorized into two subgroups: Intellectually Able (hereinafter ASD-IA; those with Wechsler’s full-scale IQ, FSIQ, and adaptive behavior composite >85, n=34) and Intellectual Impairment (hereinafter ASD-II; those with adaptive function and/or FSIQ <85, 1.5 standard deviations lower from the TDC norm; n=31). The cutoff for “Intellectual Impairment” was defined because autistic children with borderline IQ (i.e., 70-84) have similar developmental trajectories to those combined with intellectual disabilities [33]. Adaptive function was included in the definition, because intelligence alone is imprecise to predict functional abilities of autistic individuals [34]. Moreover, we intentionally did not adopt the terms “high/low-functioning” herein to preclude prejudicial and inaccurate descriptions [34, 35].

In the ASD-II subgroup, there were 24 individuals with intellectual and cognitive impairment but fair verbal expression capacity (hereinafter ASD-II-Only); 15 out of these 24 ASD-II-Only had FSIQ<70. The other 7 participants in ASD-II had an MV status (hereinafter ASD-MV; defined by the effective use of <30 words in verbal expression as reported by their parents [17])

Seven participants with ASD received ADOS Module 1; four received Module 2; twenty-four received Module 3; and thirty received Module 4. We further transformed an ADOS-2 algorithm total raw score [36] to a standardized calibrated severity score (CSS) [37, 38] to allow for the cross-module analysis.

### MRI Acquisition and Data Preprocessing

MRI data were collected on a Siemens MAGNETOM Prisma 3-T MRI with a 64-channel phased-array head/neck coil. High-resolution MPRAGE T1-weighted structural brain images were acquired sagittally using the following parameters: repetition time (TR)=2000 ms, echo time (TE)=2.43 ms, inversion time=920 ms, flip angle=9°, acquisition matrix=256×256, slice thickness=0.9 mm, in-plane resolution=0.9 mm isotropic. Multi-shell diffusion-weighted images (DWIs) were acquired using the multi-band accelerated echo-planar imaging sequence developed at CMRR [39] with the following parameters: 2.2-mm isotropic voxel, TR/TE=2238/86 ms, multi-band acceleration factor=4, number of diffusion gradient directions=19/30/90/60 at b=0/350/1000/3000 s/mm^2^, respectively. Additional b=0 images with opposing phase encoding polarity were acquired for the correction of image distortion and motion (as described below). Support procedures for participants’ MRI scans are detailed in Supplement.

DWI data pre-processing included denoising [40], Gibbs ringing removal [41], corrections for image distortions induced by eddy currents and susceptibility effects, inter-volume and slice-to-volume movements [42–44], bias field [45]; these steps were performed using MRtrix3^1^ [10] and FSL^2^ [46]. Quality assessments were performed to exclude the raw DWI data with excessive signal loss, artifacts, or in-scanner motion (based on the average root-mean-square displacements between DWI volumes, relative RMS, >1mm). The preprocessed DWI data of 38 TDC and 65 autistic participants were up-sampled and eventually analyzed (Table 1) using MRtrix3’s FBA described in the next section with recommended pipeline and parameters [10]. We also conducted complementary analysis, using the metrics FA and MD based on the diffusion tensor model, to investigate how specific the FBA results were (detailed in Supplement). The codes of the detailed preprocessing and DWI modeling are available at https://osf.io/vskj7/.

### Metrics and Statistics of FBA

The multi-shell multi-tissue constrained spherical deconvolution was applied on each upsampled DWI to compute fiber orientation distributions (FODs) of WM and tissue compartments of GM and cerebrospinal fluid [47], followed by intensity normalization to correct for compartmental inhomogeneities [48]. A study-specific FOD template was created from all TDC and ASD-Whole participants using the FOD-guided registration [49], followed by FOD segmentation [50] to generate the template fixels (FOD threshold=0.06). All participants’ fixel-wise measures were computed and mapped onto the corresponding template fixels. The fixel-wise metrics include: fiber density (FD), measuring intra-axonal volume of specific WM fiber bundles within voxels; fiber cross-section (FC), measuring the entire macroscopic/volumetric change of a fiber bundle; a combined measure of FD and FC (FDC), quantifying the overall ‘connectivity’ via microscopic density and macroscopic cross-sectional area of a fiber bundle (detailed interpretations in [12]).

We performed the statistical analysis of whole-brain fixel-wise metrics using the general linear model (GLM) incorporated with the connectivity-based fixel enhancement (CFE) approach, which implements tract-specific smoothing and thus improves test-statistics of the fixel data [51]. A whole-brain tractogram was generated using the template FODs, post-processed using spherical-deconvolution informed filtering of tractograms [50], and then used to compute fixel-to-fixel connectivity required for fixel data smoothing and enhanced statistics [51]. Two types of analyses were conducted as follows:

- Categorical analysis – As described previously, our study cohort includes two main groups (TDC and ASD-Whole) and four ASD subgroups (ASD-IA, ASD-II, ASD-II-Only, and ASD-MV) subdivided from the ASD-Whole. Whole-brain fixel-wise differences were tested between pairs of these groups, including TDC versus each ASD (sub)group, and additionally, ASD-IA versus ASD-II and ASD-II-Only versus ASD-MV. With the limited sample size, the contrast of ASD-MV with ASD-II-Only was intended for initial investigations of the effects of verbal expression capabilities but should be considered preliminary.
- Dimensional brain-behavior analysis – Mass-univariate GLMs for the whole sample (ASD-Whole plus TDC) were separately constructed to investigate which fixel (dependent variable) in the brain could be predicted by each of the following independent variables, including non-verbal IQ (NVFIQ) using the Leiter-R, adaptive behavior composite, the SRS total raw score, and global executive composite (GEC) on the BRIEF. A behavior-by-diagnosis (TDC vs. ASD) interaction term was also included in these models. Notably, we focused the brain-behavior dimensional analysis on the fluid intelligence (Leiter’s NVFIQ) because individuals in the ASD-MV subgroup cannot complete Wechsler’s verbal intelligence assessment. Another consideration was worth noting: Although the SRS manual provides T-scores for clinical screening use, the raw score is commonly used in research to estimate impairment in social functioning in people with ASD [52]. In addition, within the ASD-Whole group, we also undertook a mass-univariate GLM to investigate the brain-behavior relationship with an independent variable of ADOS-2 CSS.

For both analyses above, the nuisance variables included participant’s sex, age, medication, ICV, and relative RMS. Considering the potential age effects on behavior [53] and brain metrics [54, 55], we also additionally tested whether there were correlations between the age and aforementioned behavior/cognition variables, as well as whether there was an age diagnosis effect on FBA metrics. All these age-related tests yielded null results. The family-wise error (FWE) corrected *P*-value (hereinafter *P-FWE*) was assigned to each fixel using non-parametric permutation testing over 5,000 permutations. The outcomes for these two analyses reported in the Results section below were considered statistically significant when per-fixel *P-FWE*<0.05.

To identify the associated WM anatomy with fixels, we used TractSeg to produce labeled WM fiber bundles [56]. When an identified fixel was located close to the GM or there were no major WM tracts nearby, we applied the cerebral AAL atlas [57] and the cerebellar SUIT atlas [58] to obtain neuroanatomical labels. Yeo’s 7-network parcellation [59, 60] was used to designate the functional organization to which these fibers belong.

## RESULTS

### Demographics

As shown in Table 1, all three groups (TDC, ASD-IA and ASD-II) had comparable distributions of sex, age, relative RMS, and intracranial volume (*P*>0.05). The ratio of medication uses and co-occurring psychiatric disorders were similar between ASD-IA and ASD-II (*P*>0.05; detailed in Table S1). With significant differences in Kruskal-Wallis test (*P*<0.001), post-hoc tests showed that the ASD-IA and TDC groups were matched for intellectual function measured by Wechsler’s and Leiter’s batteries. The ASD-IA group had intermediate levels of overall adaptive function in-between the ASD-II and TDC groups. ASD-IA and ASD-II had worse executive function than TDC. ASD-IA had milder autistic symptoms based on both the SRS and ADOS-2 CSS.

### Categorical Comparisons

The ASD-Whole group, relative to the TDC, had smaller FD in the premotor segment and splenium of CC, alongside the left cerebellum Crus I (default-mode network), and smaller FDC in the right cerebellum Crus II (frontoparietal network; *P-FWE*<0.05; Figure 1, top row). Compared to the TDC group, ASD-II had smaller FD in the anterior (interconnecting bilateral prefrontal cortex) and posterior (part of the isthmus-splenium), and showed smaller FDC in the CC segment interconnecting bilateral premotor cortex, as well as in the right Crus II (frontoparietal network; *P-FWE*<0.05; Figure 1, middle row). Within the ASD-II subgroup, participants with ASD-II-Only, relative to ASD-MV, had greater FD in the middle segment and isthmus-splenium of CC (*P-FWE*<0.05). Similar to the FD result, the ASD-II-Only subgroup had greater FDC in the isthmus-splenium of CC than the ASD-MV subgroup (*P-FWE*<0.05; Figure 1, bottom row). There was no statistically significant difference in FBA metrics between the TDC and ASD-IA groups, or between the ASD-IA and ASD-II groups (*P-FWE*>0.05).

**Figure 1:**
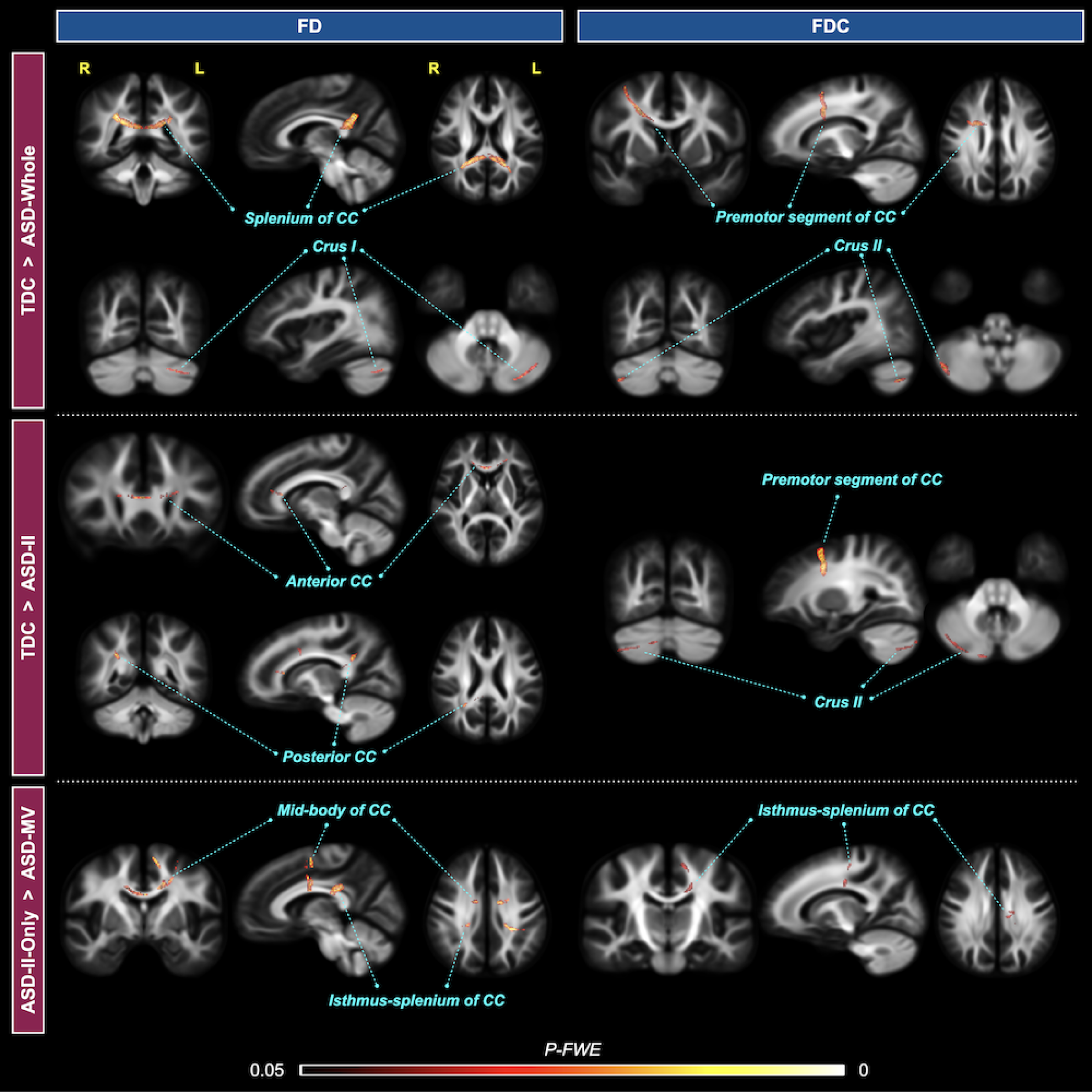
Results of categorical comparisons from the fixel-based analysis. White matter tract segments that have significant group differences in FD (left column) and FDC (right column) metrics are displayed and colored by the family-wise error corrected *P*-value (*P-FWE*). Upper block T⎯ ASD; middle blockTDC > ASD-II; bottom block⎯ ASD-II-Only > ASD-MV.

Given the spatial overlap, we extracted mean FD values from the common area of CC-nium shared between ASD-Whole vs. TDC and ASD-II vs. TDC. Figure S1 illustrates distributions of this FD which showed that ASD-IA had a higher mean and smaller standard deviation of FD (0.728±0.036) than ASD-Whole (0.725±0.043) and ASD-II (0.722±0.050), respectively. Together with its smaller sample size than that of ASD-Whole, this data feature result in null statistically significant differences between ASD-IA and TDC, alongside ASD-II, respectively (see detailed interpretations in Supplement).

### Dimensional Brain-Behavior Relationship

From the whole-sample GLMs, Figure 2 shows that NVFIQ significantly positively correlated with FC (Figure 2a-d) and FDC (Figure 2e-h) of fiber bundles in the cerebellum lobule VI (ventral attention/salience network (*P-FWE*<0.05). Significant GEC X diagnosis interactions were identified at the fibers at the GM-WM borders of the right dorsolateral prefrontal cortex (DLPFC, middle frontal gyrus subregion belonging to ventral attention/salience network; *P-FWE*<0.05; Figure 3a-b for FD, 3e-g for FDC). Specifically, GEC positively correlated with the DLPFC FD/FDC in TDC, while this brain-behavior association was negative in the ASD-Whole group (Figure 3d&h). We did not find significant correlations or diagnosis x behavior interactions of the SRS and VABS with fixel metrics (*P-FWE*>0.05).

**Figure 2:**
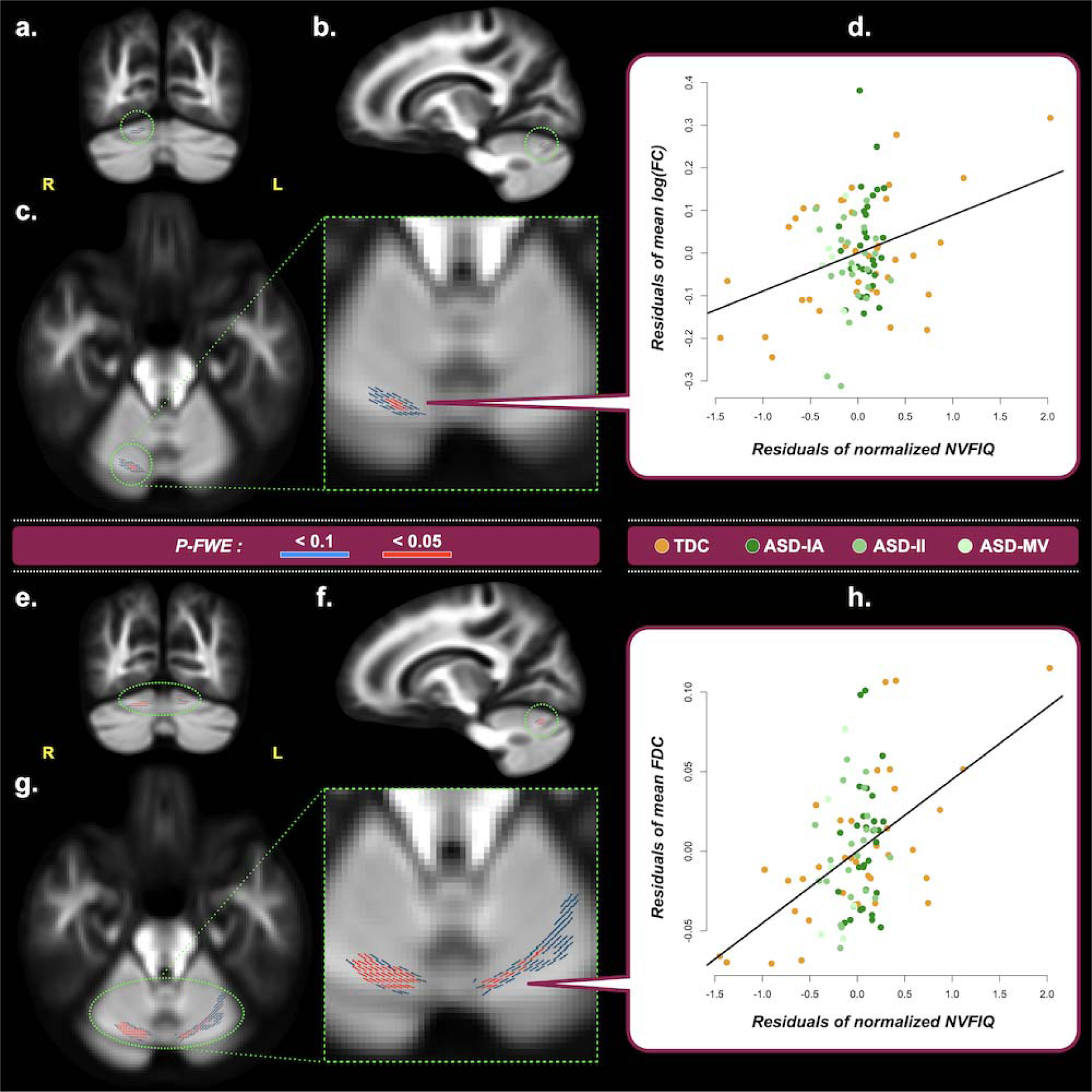
Correlations between fixel metrics and NVFIQ, derived from the dimensional analysis of the entire study cohort. Upper block ⎯ Panels a-c show the fixels where the correlation of log(FC) and Leiter’s non-verbal full-scale IQ (NVFIQ) reach statistical significance from the coronal, sagittal, and transverse view, respectively. A zoomed image of (c) is displayed with the green dashed border. Fixels are colored in red for *P-FWE*<0.05; fixels colored in blue indicate *P-FWE*<0.1 and are used to assist identification of the associated brain structure. (d) The scatter plot shows the residuals of the mean log(FC) on the vertical axis and of the normalized NVFIQ on the horizontal axis. Only all fixels that reach *P-FWE*<0.05 are considered in the plot. Lower block ⎯ The format in Panels e-h is the same as the upper block, except that the results are obtained from the analysis of FDC and NVFIQ. Acronyms ⎯ R: right; L: left; *P-FWE*: family-wise error corrected *P*-value.

**Figure 3:**
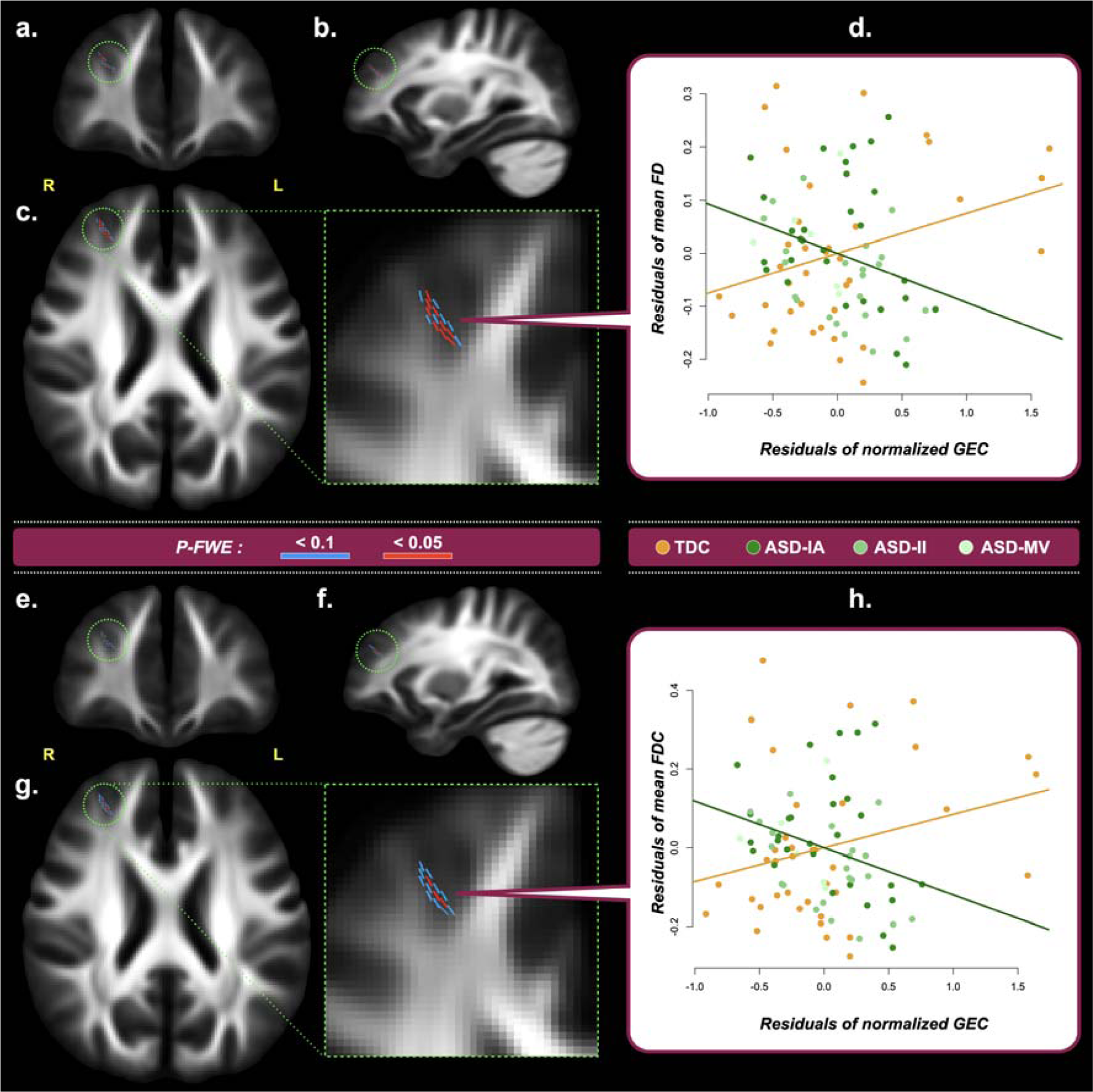
Correlations between fixel metrics and Global Executive Composite (GEC), derived from the dimensional analysis of the entire study cohort. Upper block ⎯ Panels a-c show the fixels where the correlation of FD and GEC×diagnosis interactions reach statistical significance from the coronal, sagittal, and transverse view, respectively. A zoomed image of (c) is displayed with the green dashed border. Fixels are colored in red for *P-FWE*<0.05; fixels colored in blue indicate *P-FWE*<0.1 and are used to assist identification of the associated brain structure. (d) The scatter plot shows the residuals of the mean FD on the vertical axis and of the normalized GEC on the horizontal axis. Only all fixels that reach *P-FWE*<0.05 are considered in the plot. Lower block ⎯ The format in Panels e-h is the same as the upper block, except that the results are obtained from the analysis of FDC and GEC. Acronyms ⎯ R: right; L: left; *P-FWE*: family-wise error corrected *P*-value.

The within-ASD GLM yielded that autistic individuals’ ADOS-2 CSS negatively correlated with WM FC in the right cerebellum Crus I (default-mode network; *P-FWE*<0.05; Figure 4a-d). Figure 4f shows cerebellar FA values are low (∼0.2), which cannot serve meaningful analysis. Figure 4e&g illustrate that the cerebellar FBA FODs constituted a distinct structural organization that statistical analysis can be based upon.

**Figure 4:**
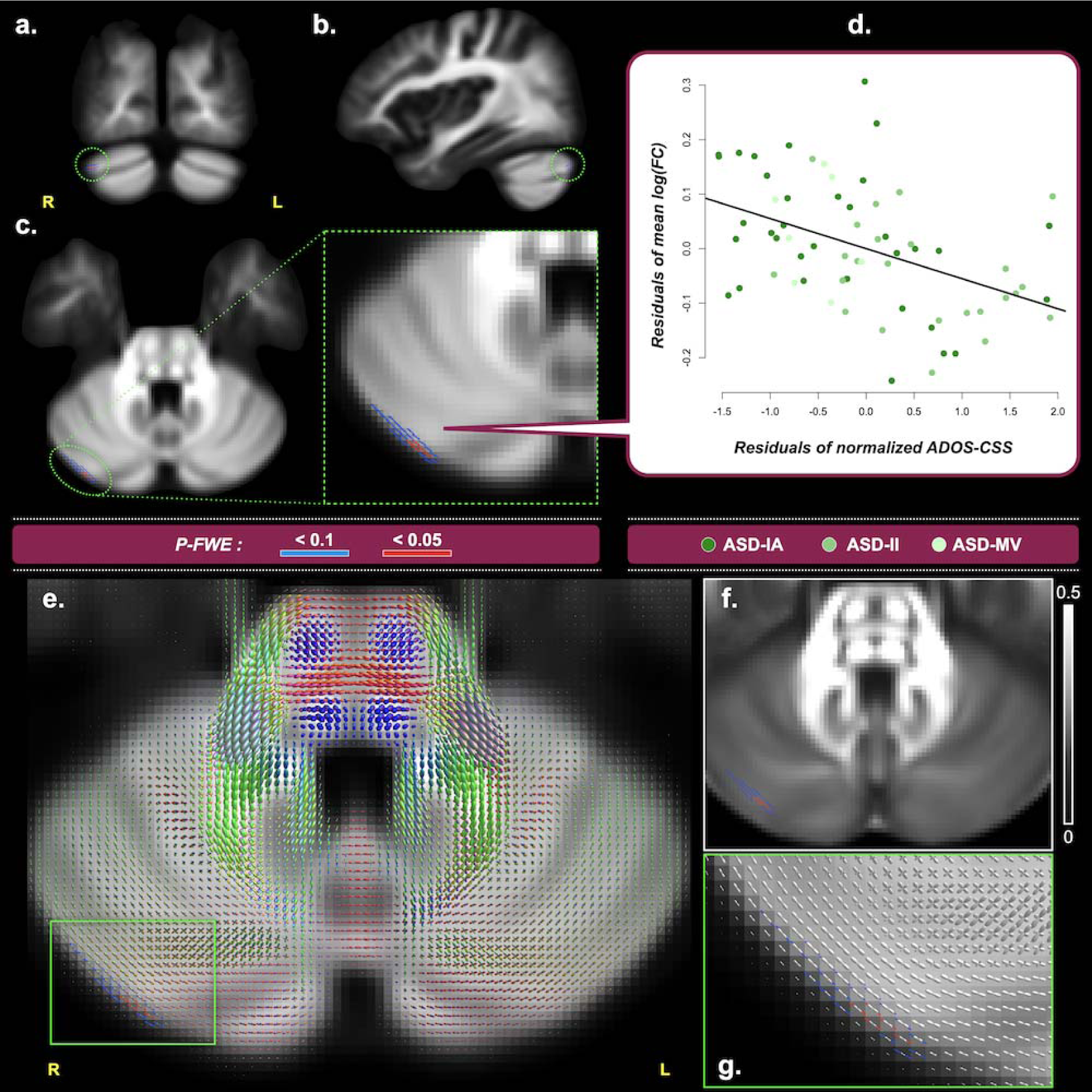
Correlations between fixel metrics and Autism Diagnostic Observation Schedule 2 Calibrated Severity Score (ADOS-CSS), derived from the dimensional analysis of the entire ASD cohort (i.e. ASD-Whole). Upper block ⎯ Panels a-c show the fixels where the correlation of log(FC) and ADOS-CSS reach statistical significance from the coronal, sagittal, and transverse view, respectively. A zoomed image of (c) is displayed with the green dashed border. Fixels are colored in red for *P-FWE*<0.05; fixels colored in blue indicate *P-FWE*<0.1 and are used to assist identification of the associated brain structure. (d) The scatter plot shows the mean log(FC) residuals on the vertical axis and the normalized ADOS-CSS residuals on the horizontal axis. Only all fixels that reach *P-FWE*<0.05 are considered in the plot. Lower block ⎯ (e) shows an axial slice of the group average FOD image. (f) shows the FA map of the same slice of (e). (g) is a zoomed region of (e), showing the microstructural organization of axonal fibers around the significant fixels within the cerebellum, which could not be reliably studied using tensor-based voxel-based analysis. Acronyms R: right; L: left; *P-FWE*: family-wise error corrected *P*-value.

### Robustness Testing

This study adopted three strategies to test the robustness and specificity of current findings. First, during the generation of the template fixels, thresholding the amplitude of the FOD lobe is required to minimize the misalignment of fixels across subjects; such an effect is particularly pronounced at the GM-WM interface where registration could be imperfect [51]. Hence, to ensure that the results were not driven by the choice of the FOD threshold, three additional FOD cutoff values of 0.04, 0.08, and 0.10 were used to construct the fixel template, and all the subsequent analyses including categorial comparisons and dimensional analysis were repeated. The main results in the categorical analysis above remained largely the same across different FOD thresholds (Figure S2). The repeated dimensional analysis also showed consistent findings in the correlated brain structures, despite some variations in fixel locations (Figure S3). Second, a more stringent significance level of *P-FWE*<0.01 was applied to test the current results. Few of the results met this strict criterion and they had reduced spatial extent of fixels (Figures S4-5). Lastly, VBA was performed with all the analyses repeated based on FA and MD. VBA neither replicated any of the results obtained from FBA nor provided additional findings (Figure S6).

## DISCUSSION

Using the current state-of-the-art dMRI sequence and FBA in a cohort with ASD across the functional spectrum, we found that ASD-associated alterations in fixel-based metrics, as identified in ASD-Whole vs. TDC, were largely driven by those combined with intellectual impairment and/or potentially by minimally verbal status (Figure S1). In addition to the altered CC, consistent with earlier literature [7, 13], we provide initial evidence suggesting that cerebellar WM also categorically and dimensionally links with the clinical presentation of ASD. Our findings highlight an essential to include autistic participants with wide-range functional levels in research, to enable a greater neurobiological understanding of the variety of functioning and cognitive profiles that manifest across the autism spectrum.

The present categorical comparison between ASD-Whole and TDC identified reduced FD in the splenium section of the CC and reduced FDC in the premotor segment of the CC, consistent with an earlier FBA report on intellectually able individuals with ASD [13]. These alterations of the CC fibers are compatible with previous dMRI [7] and other modalities [61, 62] literature suggesting impaired interhemispheric connection might contribute to idiosyncratic socio-emotional behaviors and sensorimotor integration associated with ASD. However, the altered CC microstructure appears not specific to ASD [63]. This long-standing challenge in finding disorder-specific neuroimaging biomarkers might be better disambiguated with an advanced approach of linking microscale genomic and macroscale neuroimaging observations [64].

In addition, we identified the ASD-Whole group showed altered geometries of fiber-bundle populations in the cerebellum Crus I/II. Moreover, autistic symptoms based on the ADOS correlated with the fiber bundle morphometry in the right Crus I. The left-side Crus I/II supports the socio-emotional inferences and motion-related imitation [65, 66], whereas the right counterpart subserves the complex socio-emotional reasoning and sensorimotor integration [67, 68] in neurotypical people. The GM morphometry and function of Crus I/II are reported to be involved in impaired socio-emotional [68] and language processes, alongside repetitive behaviors [67] in people [69] and mice models [70, 71] with ASD. Intellectually able youth with ASD also have specific developmental changes in GM morphometry at these loci [55, 68]. Moreover, the identified Crus I and Crus II regions are separately functionally located in the default-mode and frontoparietal networks, of which altered cerebral activity and connectivity have been observed in ASD [72, 73]. Taken together, consistent with the functional MRI literature, we provide the first empirical evidence to imply that Crus I/II WM essentially links with autistic conditions. However, whether cerebellar lateralization plays a role in ASD-associated fiber pathology awaits further confirmation.

When comparing ASD-IA with TDC, with the similar sample size and demographic features to the participants of Dimond *et al.*’s work [13], we did not replicate the significant ASD-associated WM differences. Conversely, the ASD-II vs. TDC differences were similar to ASD-Whole vs. TDC differences in the CC-splenium (based on FD) and right Crus II (based on FDC). Despite the preliminary nature due to a small sample size of ASD-MV, the difference between ASD-II-Only and ASD-MV also was located at the posterior CC. As shown in Figure S1, our findings suggest that ASD-associated alterations in the geometry of fiber-bundle populations, as identified by ASD-Whole vs. TDC, are mainly driven by the fiber pathology in those autistic people with lower intellectual/adaptive function. This uncovering might help reconcile parts of inconsistency in the previous FBA [13, 14] and other MRI literature [74]. This is because these neuroimaging studies chiefly based on intellectually able individuals [18] tend to show smaller effect sizes [75] and be susceptible to sampling heterogeneity [76]. Our results also highlight the imperativeness of having research practices be inclusive of all autistic individuals on the spectrum, especially the understudied population with developmental disabilities [19, 20], in order to address a gap of better characterizing these diverse autistic samples and development of more individualistic targeted plan accordingly [17, 18, 21].

The dimensional brain-behavior analysis showed that the positive correlation between fluid intelligence and FC/FDC metrics in the cerebellum lobe VI existed cutting across ASD and TDC as a continuum. The higher the non-verbal intelligence is, the stronger the fiber-bundle populations are. Earlier functional MRI studies reported that the cerebellar VI is involved in motor processing, and nonmotor cognitive processing is implicated in language and working memory in ASD [77–80]. This posterolateral cerebellar area also supports the foundational function of hierarchical cognitive control such as mental arithmetic [81]. This lobule also participates in the ventral attention/salience network [82], whose cerebral connectivity links with neurotypical individuals’ fluid reasoning capacity [83]. Taken together, the current result suggests the essential role of the posterior cerebellum in contributing to human fluid cognition, regardless of the neurodiverse status.

Moreover, we identified a diagnosis-specific relationship between the summarized measure of daily-life executive function and the FC/FDC in the GM-WM borders of the right DLPFC in the ventral attention/salience network. Individuals with ASD usually have a broad executive dysfunction that is relatively stable across development [84]. The DLPFC is one of the primary areas that support executive function processing such as planning and working memory [85, 86]. This prefrontal area plays an essential role in executive dysfunction associated with ASD across ages [87]. Moreover, ventral attention/salience network appears to link with ASD [88] and self-control [89] in different directions. ASD has poor delineation of the GM-WM boundary at the DLPFC [90, 91]. Collectively, while future studies are required to establish the relationship between the fiber-bundle populations and the differentiation of the GM-WM boundary, it is likely that altered neurodevelopmental processes involving WM fibers at the GM-WM boundary contribute to executive dysfunction commonly observed in ASD.

The capability of FBA to quantify fiber-specific measures of micro-/macro-structural alterations has been shown to be more beneficial for cerebral WM over VBA [11, 92]. Advancing previous findings based on the DTI model [19, 20] and advanced model of structural T1 image [90], our study reveals that the cerebellum and the WM underneath as well as the GM-WM junction at the DLPFC are involved in ASD pathology and associated cognition. The supplementary analysis (Figure S1-4) not only supports the robustness of these findings, but also suggests that provided the equivalence with the present dMRI data quality in combination with multi-tissue modeling [47], FBA could be advantageous to detect small-yet functionally meaningful alterations in voxels containing complex arrangement of axonal compartments, such as at the GM-WM partial volume voxels or cerebral/cerebellar parenchyma. Notably, these findings were only observed using FBA, whereas VBA based on diffusion tensor measures did not yield any significant results. This discrepancy may originate from that tensor-based metrics are prone to the effects of crossing fibers and therefore more susceptible to noise contamination at those brain regions [93]. Among other explanations of sampling heterogeneity [76] and head motion [94], the current strength of using advanced analysis technique might also partly account for the null finding in the uncinate fasciculus, inconsistent with earlier meta- [5] and mega-analyses [63]. This WM tract is especially liable to false-positive findings because of the crossing fibers [95, 96].

### LIMITATIONS

First, the sample had a relatively wide age range. Although our additional age-related tests yielded null results, we cannot exclude residual developmental effects on the current findings. Second, the present sample was male predominant, limiting any formal tests on sex-related differences [97]. Third, we recruited participants from outpatient clinics, with many showing psychiatric comorbidities commonly associated with ASD (such as ADHD and anxiety disorders) or/and psychotropic medication uses. On the flip side, this design may render the current results more generalizable to ‘real-world autism’, as these psychiatric comorbidities may affect up to 70% of individuals with ASD [98]. The intellectually able participants with ASD without (or with restricted proportions of) psychiatric comorbidities in most of earlier neuroimaging literature actually belong to an outlier subgroup on the spectrum [76]. Moreover, there was no difference in ratios of comorbidity and medication utilization between the ASD subgroups, which reduces the doubt that the current results might be driven by such comorbidities. Fourth, our sample size was relatively small, limiting statistical power to detect small between-group effects and paradoxically increasing the risk of inflating effect sizes [99]. In particular, the outcomes related to ASD-MV should be considered preliminary. Lastly, we did not include intellectual impairment-only without ASD as an additional control condition. Scientific progress will benefit from intellectually-informed recruitment strategies to discover where ASD and intellectual impairment intersect and part ways.

## CONCLUSION

In sum, we confirmed that changes in anatomical connections linking hemispheres, as well as cerebellum Crus I/II might contribute to autism phenotypes. These ASD-associated alterations appear to be mainly driven by autistic individuals with intellectual impairment and potentially further by minimally verbal status. Across the functional spectrum, autistic severity, nonverbal intelligence and executive function associated with WM fiber-bundle properties in regions where WM pathology based on dMRI are seldom reported. These results highlight that by embracing the inclusion of understudied sub-populations on the spectrum, together with the development of novel neuroimaging methods, we may better reconcile heterogeneity across studies and advance the understanding of the neuropathology of ASD.

## List of Abbreviations

ADI-R: Autism Diagnostic Interview-Revised
ADOS: Autism Diagnostic Observation Schedule
ADOS-CSS: Autism Diagnostic Observation Schedule 2 Calibrated Severity Score
ASD: autism spectrum disorder
BRIEF: Behavior Rating Inventory of Executive Function
CC: corpus callosum
DLPFC: dorsolateral prefrontal cortex
dMRI: diffusion magnetic resonance imaging
DWI: diffusion-weighted image
FA: fractional anisotropy
FBA: fixel-based analysis
FD: fiber density
FC: fiber cross-section
FDC: fiber density and cross-section
FOD: fiber orientation distribution
FSIQ: Full-Scale Intelligence Quotient
GEC: global executive composite
GLM: general linear model
GM: gray matter
IA: intellectually able
ICV: Intra-Cranial Volume
ID: intellectual disabilities
II: intellectual impairment
K-SADS-E: Kiddie-Schedule for Affective Disorders and Schizophrenia-Epidemiological Version
Leiter-R: Leiter International Performance Scale-Revised
MD: mean diffusivity
MV: minimally verbal
NVFIQ: Non-Verbal Intelligence Quotient
P-FWE: family-wise error corrected p-value
RMS: root-mean-square
SRS: Social Responsiveness Scales
TDC: typically developing controls
VBA: voxel-based analysis
VABS: Vineland Adaptive Behavior Scales
WAIS-IV: Wechsler Adult Intelligence Scale-4th edition
WISC-IV: Wechsler Intelligence Scale for Children-4th edition
WM: white matter

## ETHICS DECLARATION

### Ethical approval and consent to participate

The study was approved by National Taiwan University Hospital, Taiwan, and written informed consent was obtained from all participants or their legal guardians (for participants < 18 years).

### Availability of data and material

The data that support the findings of this study are available from the corresponding author, H-YL, upon reasonable request.

### Consent for publication

Not applicable.

### Authors’ contributions

All named authors contributed to the manuscript. H-YL designed the experiment and led the project. C-HY, R-YT and H-YL analyzed and interpreted the data and drafted the manuscript. H-CN, LC and SSG interpreted the data. T-LC and SSG developed the stimuli and measures. H-YL, H-CN, Y-YW and SSG were responsible for the collection of the clinical data. H-YL, M-YH, J-CC and E-NT contributed to neuroimaging data collection. H-YL, C-HY, T-LC and SSG acquired funding. All authors read and approved the final manuscript.

## Acknowledgements

The authors would like to thank all of our participants and their family members for partaking in this study, Ms. Yi-Chun Liu for carrying out the recruitment and data management, Imaging Center for Integrated Body, Mind and Culture Research, NTU for equipment support, and the anonymous reviewers for comments that significantly improved the manuscript.

## Supplementary Material

### A.

**Supplementary Table S1.** Comorbidity and Medication Use for ASD-IA & ASD-II.

A. Supplementary Table S1. Comorbidity and Medication Use for ASD-IA & ASD-II
|  | ASD-IA (n= 34) | ASD-II (n= 31) |
| --- | --- | --- |
| **Comorbidity** |  |  |
| No comorbidity | 8 | 7 |
| Comorbid ADHD | 20 | 18 |
| Comorbid Anxiety disorder | 8 (5 social anxiety disorder; 1 history of mild agoraphobia; 1 height phobia) | 9 (4 specific phobia; 4 social anxiety disorder; 1 selective mutism) |
| Comorbid tic disorder | 3 | 5 (2 Tourette syndrome) |
| Comorbid OCD | 1 | 1 |
| Comorbid learning disorder | 1 (writing/reading disorder) | N/A |
| Comorbid depressive disorder | 3 (1 history of depression; 1 dysthymic disorder & mild MDD) | N/A |
| Other comorbidities | 1 gender dysphoria<br>1 ODD | 2 epilepsy (well controlled)<br>1 ODD |
| **Medications** |  |  |
| Methylphenidate | 13 | 6 |
| Antidepressant | 2 (1 sertraline; 1 fluoxetine) | 1 (sertraline) |
| Valproic acid (mood stabilizer) | N/A | 2 |
| Antipsychotic | N/A | N/A |

Acronym – ADHD: Attention Deficit/Hyperactivity Disorder; MDD: Major Depressive Disorder; OCD:
Obsessive Compulsive Disorder; ODD: Oppositional Defiant Disorder.

### B. Support procedures for MRI scans

Except the screening survey in clinics, our support strategies for MRI scans were devised and adapted from the published protocol [100]. Specifically, the clinicians in outpatient clinics made a preliminary evaluation to survey whether candidate participants with ASD (especially those with II) might be able to comply with MRI procedures. Following the first-step screening and behavioral assessments, we then shared video and recordings of scanner sounds with each participant (who shows the potential for completing the scans) and their parents two weeks before the scans, in order to help them simulate the scanning environment at home. We also offered an optional one-hour session in a mock scanner within one week before the scan day, if participants or their parents requested or desired the practice.

On the scan day, we would schedule participants 30 minutes to one hour before the formal procedure to help them reduce anxiety by slow exposure to the scan environment, including the room, bed, head coils, and scanner. All participants wore their own MRI-safe clothes without changing into scrubs. After MRI safety checks, their parents, therapist, or our research assistant were allowed to accompany and physically contact participants (touching/patting abdomen or legs or holding hands) during the scanning. This physical company was requested for every participant in the ASD-II subgroup and was optional for the others. Participants’ comfort items were also allowed in the scan room if passing safety checks. Blankets, passive noise-canceling earplugs, soft pads were prepared and opted for individual’s special sensory needs. A movie paradigm, Inscapes, which has shown efficacy to improve compliance in MRI [101], was repeatedly played during T1 and DWI acquisition, while participants were also allowed to freely close their eyes if they disliked the videoclips. We would communicate verbally with the parent of participant with ASD-II or with the participant (with ASD-IA or controls) via the microphone from the control room to ensure their physical and psychological status was good enough for the scans.

### C. Mean fiber density of the splenium of corpus callosum across groups

The outcomes of the categorical analysis shown in Figure 1 demonstrated that the significant differences of FBA metrics were found at: TDC > ASD-Whole, TDC > ASD-II, and ASD-II-Only > ASD-MV. When knowing TDC > ASD-Whole, one would expect that TDC should also have greater fixel metrics (e.g. FD) than any ASD subgroups. Thus, the results of no significant differences between TDC and ASD-IA might seem to be contradictory to such expectations. The purpose of this section is to attempt to explain the rationale behind the relevant findings.

Figure S1 shows the complementary outcomes to Figure 1, from which a mask containing “significant fixels” at the splenium of the corpus callosum was derived first. For each (sub)group, the mean and standard deviation of the FD metric within the fixel mask was then computed for each participant, yielding the group distribution of FD. Consistent with Figure 1, TDC showed significantly higher mean FD than ASD-Whole. In addition, divided from ASD-Whole, the results suggest that the distribution of ASD-IA “shifted” toward TDC, thus yielding non-significant differences between TDC and ASD-IA.

Likewise, Figure S1 also explains why there were no differences between ASD-IA and ASD-II: When separated from ASD-Whole, the distribution of ASD-II indeed “shifted” oppositely with ASD-IA. However, although both subgroups had significant differences in their clinical features (Table 1), the alteration in mean FD did not reach the statistical significance. Nevertheless, such a shift toward reduced mean FD resulted in a significantly lower value of ASD-II as compared to TDC, which was consistent with the outcomes of FBA shown in Figure 1.

Combining Figures 1 and S1 therefore leads us to the conclusion that fiber-specific alterations are driven by those combined with II and/or MV, and that it is imperative to include these populations in ASD research to avoid the selection bias.

**Figure S1:**
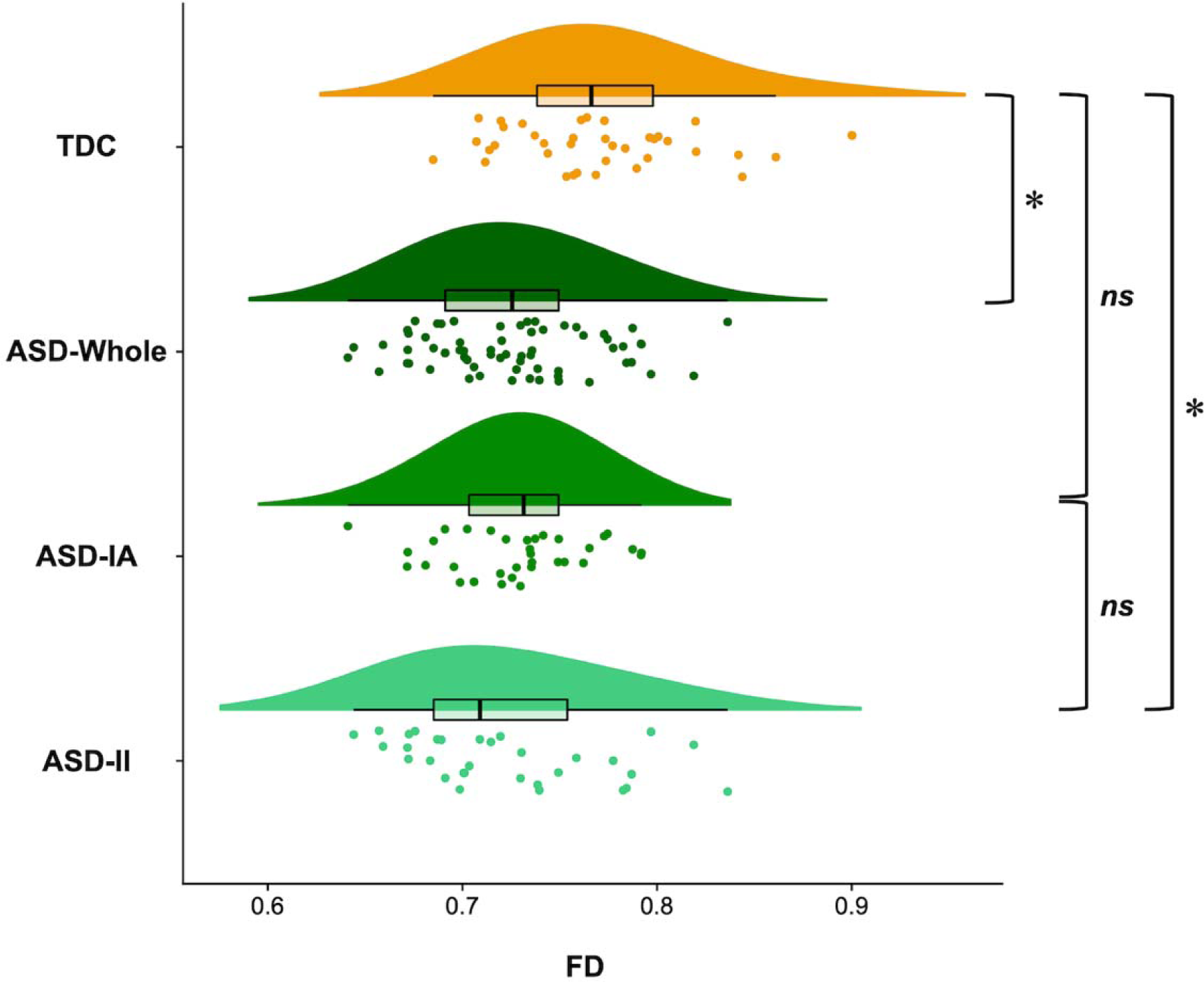
Raincloud plots showing the distribution of per participant’s mean fiber density (FD) metric in each (sub)group. FD values were extracted from fixels that showed significant reductions in FD at the splenium of the corpus callosum, which was defined by the spatial overlap between the outcome of TDC vs. ASD-Whole and TDC vs. ASD-II. “*” indicates significant differences of the distribution; “*ns*” denotes that the differences between groups are non-significant.

### D. FBA at different thresholds of FOD peak amplitudes

The threshold of the FOD peak value defines the minimum amplitude of an FOD lobe to be regarded as a fixel. Hence, lowering the threshold will include small fixels potentially representing a small fraction of fibers presenting in a voxel, at the price of increasing false positive fixels that could come from the effect of noise contamination. Likewise, increasing the threshold will make the FBA focus only on large fixels or major WM fiber bundles, it would however have a detrimental effect as the contribution of fixels below the threshold to the DWI signal will be completely discarded. The default FOD cutoff in *MRtrix3* was set to 0.06 empirically, we therefore provide the results of complementary FBA in this section, in which we adjust the value to 0.04 / 0.08 / 0.10 to ensure that the findings in the current study were not determined by the choice of the threshold. Overall, the FBA outcomes remain largely consistent following the adjustment of the FOD cutoff value. Despite some expected variations in the spatial extent of the significant fixels, no additional anatomical regions were identified.

Figure S2 shows the results of categorical comparisons under the FOD cutoff values of 0.04 / 0.08 / 0.10. Comparing with the default threshold of 0.06 shown in Figure 1, only one brain region where the group differences became non-significant: In TDC versus ASD-II, no significant difference in FD at the posterior CC was detected under the cutoff value of 0.10.

**Figure S2:**
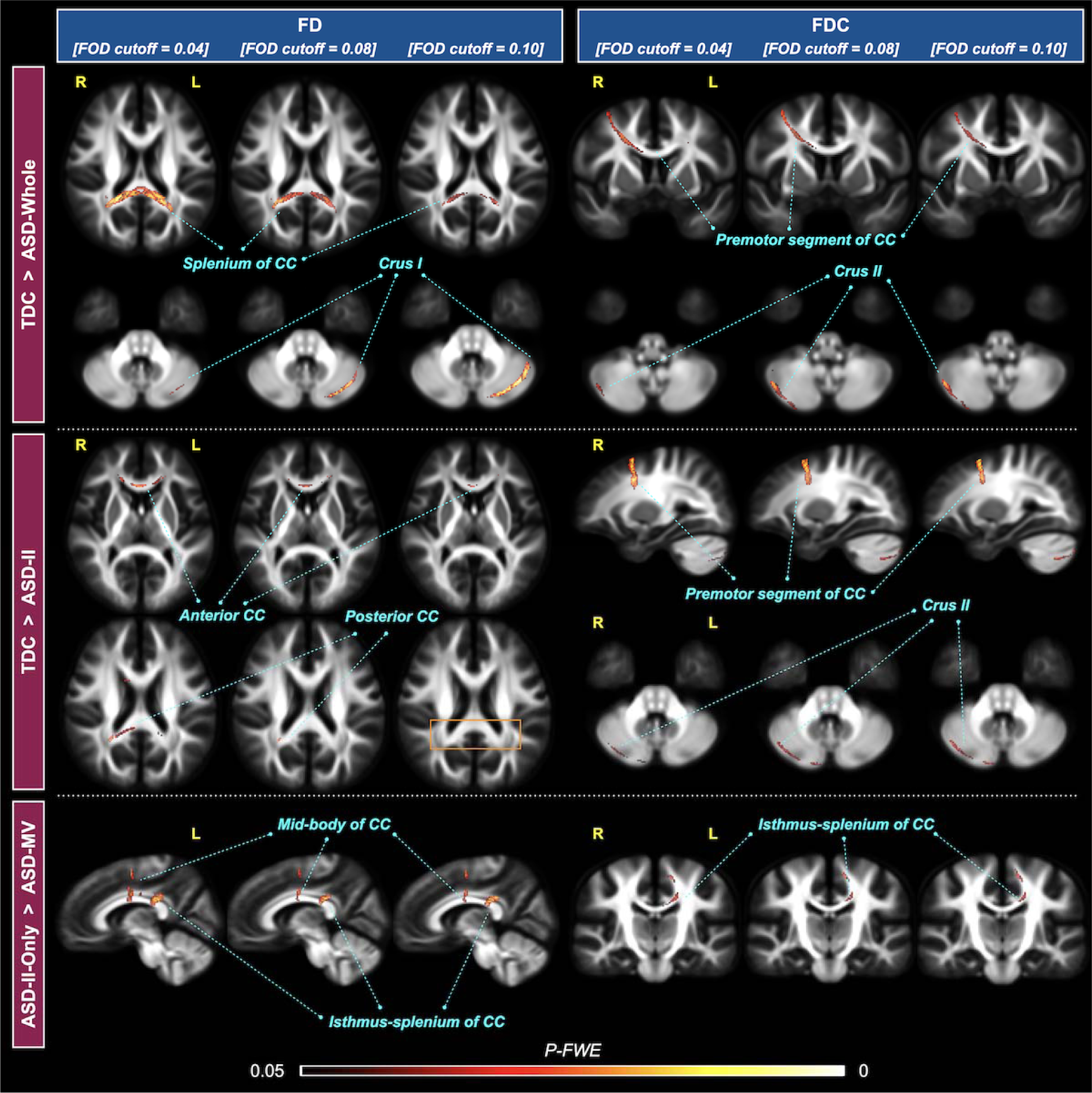
Results of categorical comparisons using an FOD cutoff threshold of 0.04, 0.08, and 0.10. Different thresholds were listed from left to right, for the FD (left column) and FDC (right column) metric. Significant fixels are colored by P-FWE and overlaid on the FOD template. Upper block, TDC > ASD-Whole; middle block, TDC > ASD-II; bottom block, ASD-II-Only > ASD-MV. The rectangle colored in orange indicates the missing significance comparing to the original results under the FOD cutoff of 0.06 in Figure 1 of the main content.

Figure S3 shows that the dimensional brain-behavior analysis under different FOD cutoff values has a similar pattern with the results shown in Figures 2 and 3.

**Figure S3:**
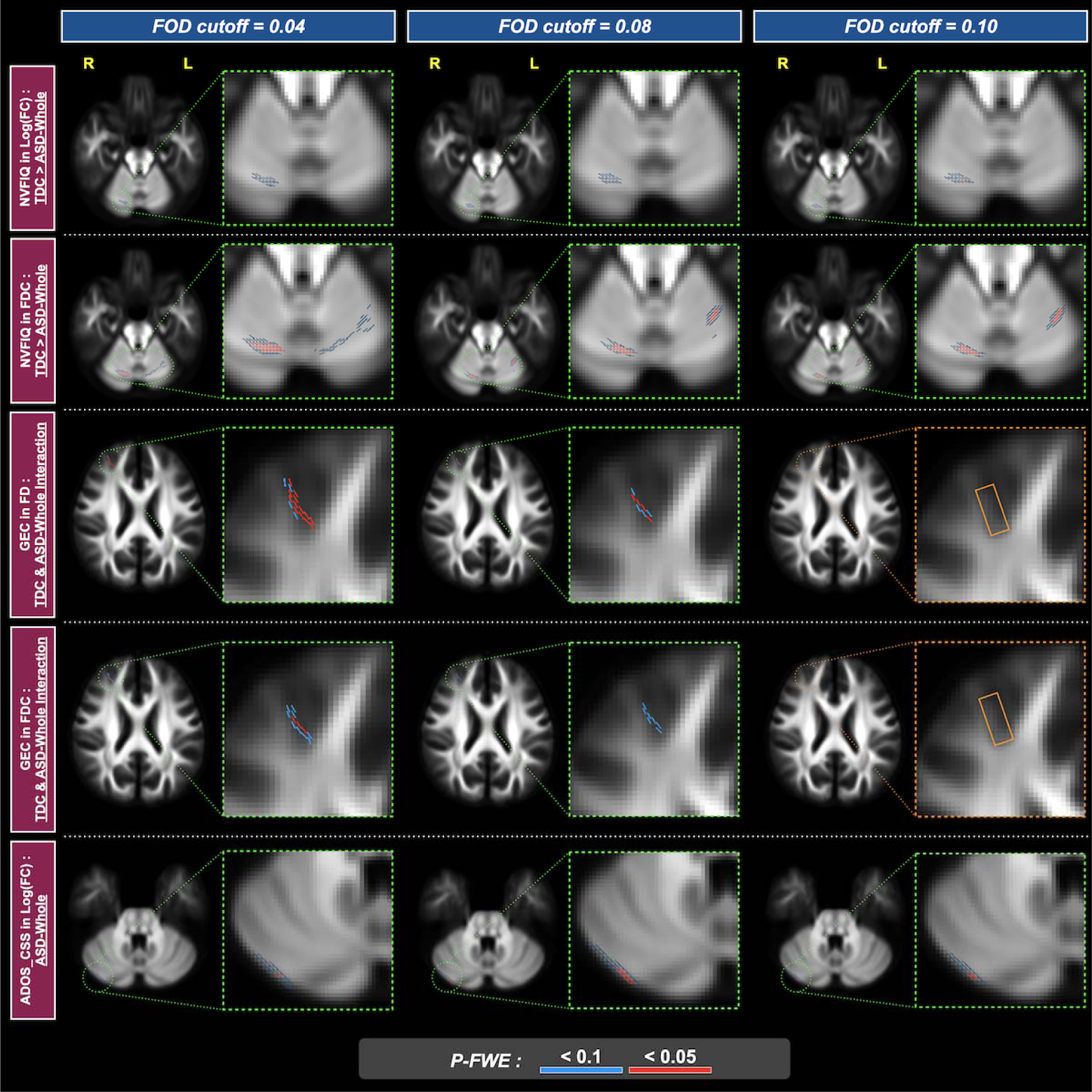
Results of dimensional brain-behavior analysis using an FOD cutoff threshold of 0.04 (left column), 0.08 (middle column), and 0.10 (right column). Each row illustrates the correlations between a fixel metric and a behavior score, in comparison to the outcomes under the FOD cutoff of 0.06 shown in Figures 2 and 3 of the main content. The rectangle colored in orange indicates the missing significance comparing to Figure 3. Fixels are colored in red for *P-FWE* < 0.05; fixels colored in blue indicate *P-FWE* < 0.1 and are used to assist identification of the associated brain structure.

Interestingly, changing the FOD threshold appears to alter the FBA results oppositely in the cerebrum and cerebellum, as observed from both the categorical comparisons and dimensional analysis above. The application of a higher FOD cutoff value results in a reduced significance level in the cerebrum but an enhanced statistics in the cerebellum, either based on the FD or FDC metric. The explanation for this contrary tendency is not straightforward, as such a tendency might be dependent on the spatial location and study-specific. Ideally, one could consider using a lower FOD threshold, given sufficient quality of dMRI data. The outcomes provided in this section suggest that the default FOD cutoff of 0.06 used in this study might be the right balance for investigating both the cerebrum and cerebellum.

### E. FBA at a significance level of *P-FWE* = 0.01

This section provides the results of using a more stringent statistical level of *P-FWE* < 0.01, with all other processing procedures and parameters identical to those in the main article. Supplementary Figures S4-5 show the results for the categorical comparisons and dimensional analysis, respectively.

**Figure S4:**
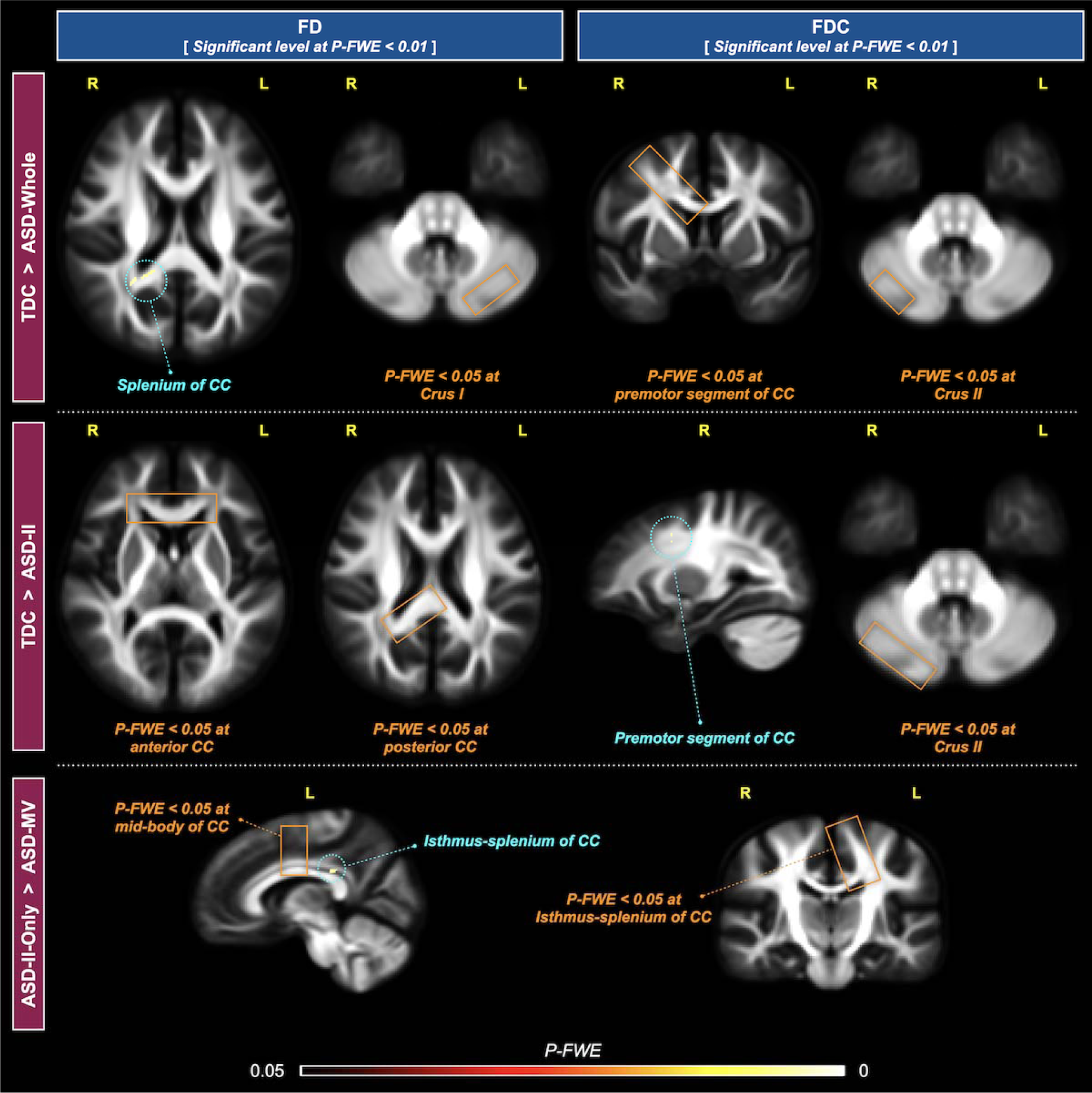
Results for categorical comparisons under a statistical threshold of *P-FWE* < 0.01. Left column – metric FD; right column – metric FDC. Upper block ⎯ TDC > ASD-Whole; middle block ⎯ TDC > ASD-II; bottom block ⎯ ASD-II-Only > ASD-MV. Fixels within blue dashed circles reached the significance level of *P-FWE* < 0.01. The rectangles colored in orange highlight brain regions where fixels reached *P-FWE* < 0.05 but did not pass the statistical testing using *P-FWE* < 0.01.

**Figure S5:**
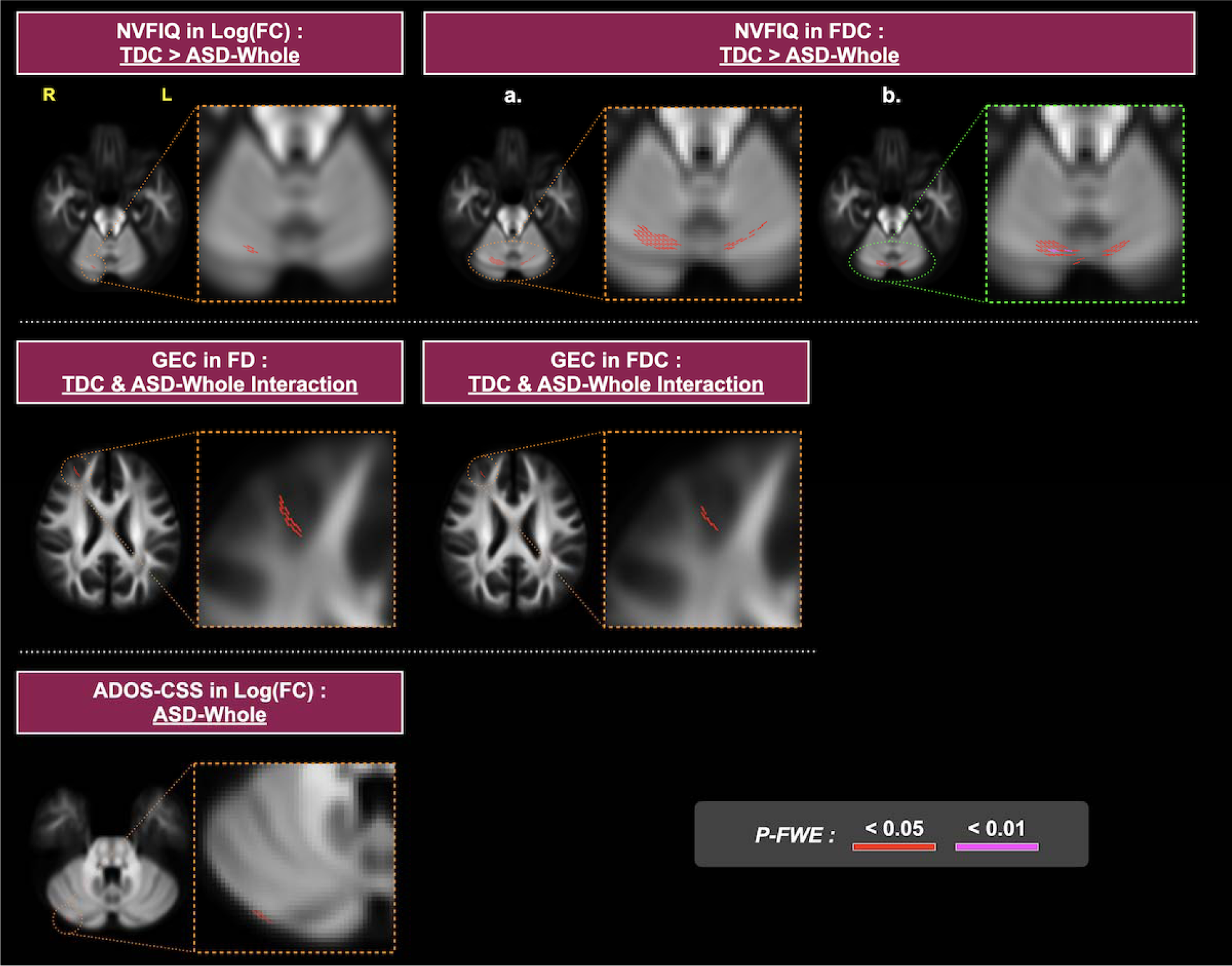
Results of dimensional analysis under a statistical threshold of *P-FWE < 0.01*. Upper row – NVFIQ: The only significance under this more restricted threshold is in the positive correlation of NVFIQ and FDC. Panel a shows the same slice in main content, where no significant results appear at *P-FWE* < 0.01. Panel b is the result at one slice below Panel a, where the significance under *P-FWE* < 0.01 are observed. Middle row – GEC: no fixels reach *P-FWE* < 0.01. Bottom row – ADOS-CSS no fixels reach *P-FWE* < 0.01. Fixels are colored in magenta for *P-FWE* < 0.01; fixels colored in red indicate *P-FWE* < 0.05 and are used to assist identification of the associated brain structure.

### F. Voxel-based analysis based on diffusion tensor model

This section provides the results obtained from the same study cohort using the conventional voxel-based analysis (VBA) based on the diffusion tensor metrics to benchmark against the present FBA outcomes. To this end, FA and MD maps were computed for each participant using the b=0 and b=1000 DWI volumes (90 diffusion gradient directions) extracted from the multi-shell DWI data at the preprocessed level. The FA maps of all TDC and ASD-Whole participants were used to generate a study-specific FA template. The FA and MD maps of each participant were then transformed to the template space via applying the transformation field obtained from the image registration between individual’s FA map and the template. A Gaussian kernel of 6-mm full width at half-maximum in size was applied to smooth the transformed FA and MD maps. The whole-brain statistical analysis was performed using GLM, with the threshold-free cluster enhancement applied. The nuisance variables included participant’s sex, age, medication, and relative RMS. All analyses including both the categorical and dimensional analysis as in FBA were performed for VBA. Nonparametric testing was performed using 5,000 permutations, and the significance level was defined at *P-FWE* < 0.05. All the processing steps above were done using *MRtrix3*.

Supplementary Figure S6 shows the voxels/regions that reached statistical significance (*P-FWE* < 0.05). For the categorical comparisons, the TDC group has higher FA than the ASD-Whole at a cluster of voxels within the third ventricle. The dimensional brain-behavior analysis showed that there was a positive correlation between FA and GEC at a cluster that was also located in the brain ventricle. Both results however cannot draw a meaningful interpretation. All the results reported from FBA in the present study were not found using either FA- or MD-based VBA.

**Figure S6:**
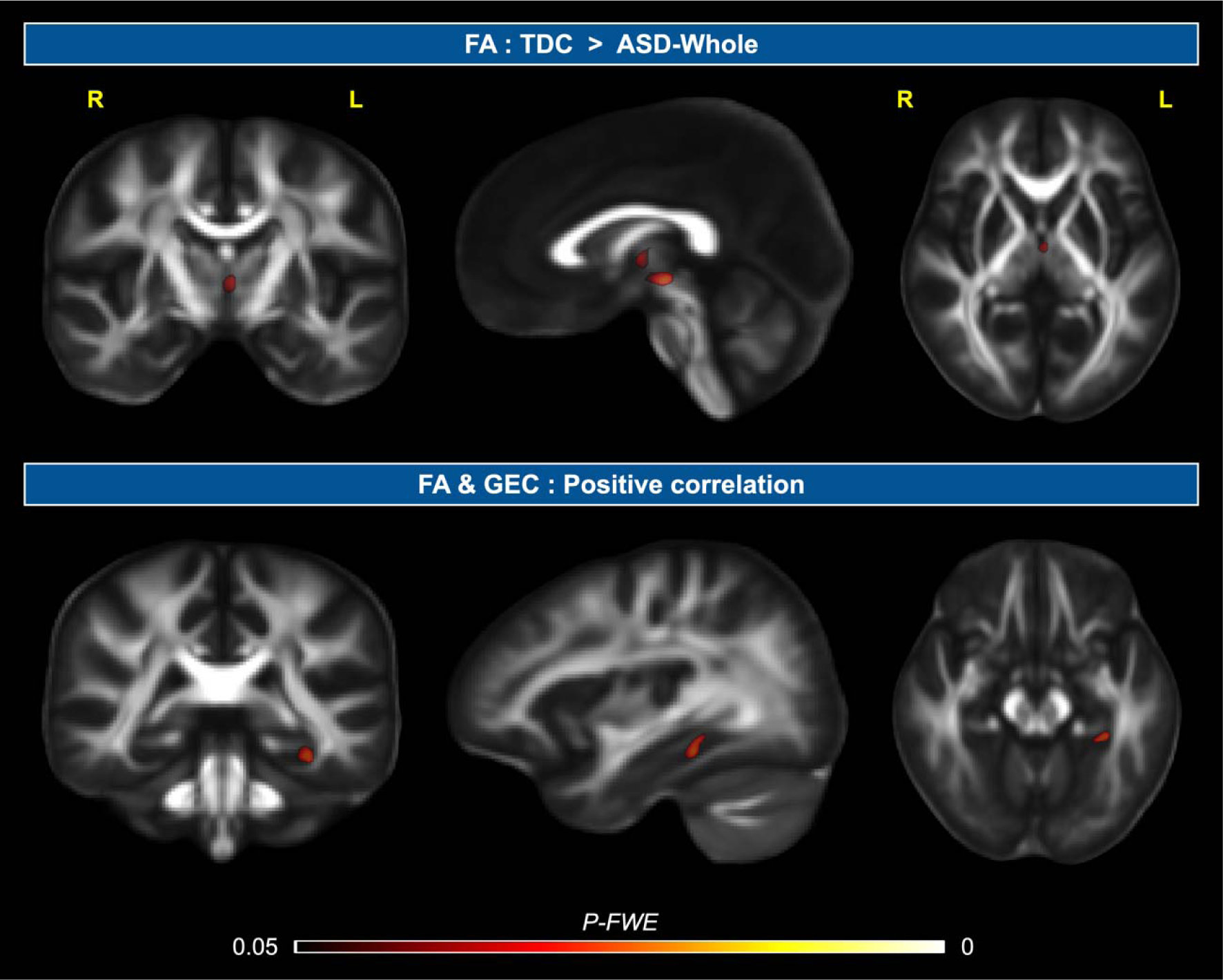
Results of VBA using the FA and MD metrics of the diffusion tensor model. Voxels that reached statistical significance (*P-FWE* < 0.05) are colored by the *P-FWE* and overlaid on the group FA template images in three orthogonal planes. Upper row ⎯ the cluster where TDC have greater FA than ASD-Whole. Bottom row ⎯ the cluster where GEC positively correlates with FA.

## Footnotes

1 https://www.mrtrix.org/

2 https://fsl.fmrib.ox.ac.uk/fsl/fslwiki/FSL

